# A novel decellularization method to produce brain scaffolds

**DOI:** 10.1101/680702

**Authors:** Alessandro E.C. Granato, Edgar Ferreira da Cruz, Dorival Mendes Rodrigues-Junior, Amanda Cristina Mosini, Henning Ulrich, Arquimedes Cheffer, Marimelia Porcionatto

## Abstract

Scaffolds composed of extracellular matrix (ECM) can assist tissue remodeling and repair following injury. The ECM is a complex biomaterial composed of proteins, glycoproteins, proteoglycans, and glycosaminoglycans, secreted by cells. The ECM contains fundamental biological cues that modulate cell behavior and serves as a structural scaffold for cell adhesion and growth. For clinical applications, where immune rejection is a constraint, ECM can be processed using decellularization methods intended to remove cells and donor antigens from tissue or organs, while preserving native biological cues essential for cell growth and differentiation. Recent studies show bioengineered organs composed by a combination of a diversity of materials and stem cells as a possibility of new therapeutic strategies to treat diseases that affect different tissues and organs, including the central nervous system (CNS). Nevertheless, the methodologies currently described for brain decellularization involve the use of several chemical reagents with many steps that ultimately limit the process of organ or tissue recellularization. Here, we describe for the first time a fast and straightforward method for complete decellularization of mice brain by the combination of rapid freezing and thawing following the use of only one detergent (Sodium dodecyl sulfate (SDS)). Our data show that using the protocol we describe here the brain can be entirely decellularized, while still maintaining ECM components that are essential for cell survival and repopulation of the scaffold. Our results also show the repopulation of the decellularized brain matrix with Neuro2a cells, that were identified by immunohistochemistry in their undifferentiated form. We conclude that this novel and simple method for brain decellularization can be used as a biocompatible scaffold for cell repopulation.

**Impact Statement:** For the first time we describe an easy, effective and low cost method for complete decellularization of murine brain by the use of only one detergent (SDS) combined with rapid freezing and thawing, that can be used as a 3D scaffold for cell culture of neuronal cells. The results show that the decellularized brains still maintain ECM components essential for cell survival and repopulation of the scaffold. Moreover, we found that the decellularized brain matrix can be repopulated with neural cells, showing its biocompatibility.

**GRAFICAL ABSTRACT:** 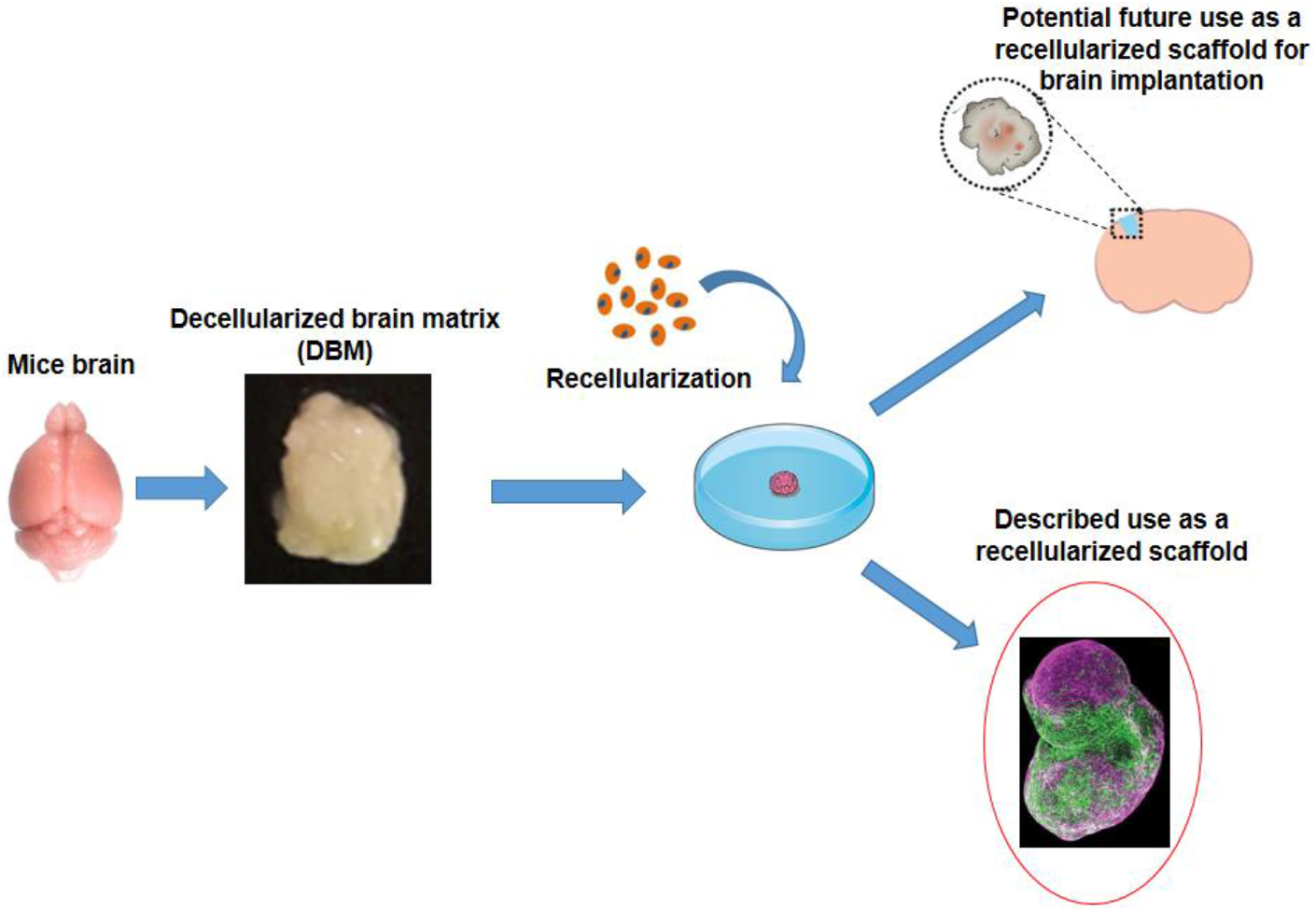

## INTRODUCTION

Chronic diseases represent a burden to the individuals as well as to the health systems worldwide. Recently, several strategies have been developed to better understand and eventually treat chronic diseases. These strategies comprehend the establishment of organoids produced from human induced pluripotent stem cells (hiPSC), bioengineered tissues, and decellularized scaffolds from lungs, hearts, livers, and kidneys ^(1–5)^. Despite the great advances till now, none of these processes can reproduce with high fidelity the full complexity of a tissue or organ. Combinations of organoids and organ-on-a-chip technology have been proposed, but still face technical difficulties, including the correct assemble of a given tissue microenvironment ^6^. Thus, one of the most challenging aspects of tissue engineering is to reproduce the correct disposition of the extracellular matrix (ECM), known not only to support correct cell biology processes, but in many cases, to drive them. The brain is an emblematic example of this issue and the production of brain organoids from hiPSC has been described in the literature by several laboratories ^(7–10)^. Interestingly, the organoids present gene expression that recapitulates human cortical development, indicating that the maturation of a human organoid runs in synchrony with healthy human fetal development ^11^. Despite the power of organoid cultures, there are some limitations such as the lack of full organ organization and orientation. Furthermore, there is a size limitation to organoids as they are not vascularized in vitro ^(12,13)^.

The solution may lay in the reproduction of the ECM composition, not only with regard to the types of proteins, glycoproteins, and proteoglycans, but also to their arrangement in the tissue ^(14–21)^. Decellularization of organs can produce ECM scaffolds that can be used for stem cell differentiation as an alternative to organoids. The ECM acts as a support for cells during embryo development, induces cell differentiation and proliferation, promotes cell adhesion and cell migration, and provides cues to maintain tissue homeostasis and regeneration. Similar to organoids, acellular organ substrates combined with patient-derived stem cells can create a platform for personalized medicine studies ^13^. Properly decellularized organs retain both the structural proteins and biologically active growth factors that are capable of inducing location-specific differentiation of stem and progenitor cells ^(13,22,23)^.

Tridimensional (3D)-printed scaffolds are an attractive alternative to decellularized scaffolds. However, it is currently difficult to use this technology to replicate the intricate ECM structure of complex organs, such as the brain, and the composition of region-specific growth factors on the ECM scaffold ^13^. To produce a bioscaffold, several steps are required in order to achieve complete decellularization and that can be performed by flushing detergents or enzymes through to the main artery of the organ. Based on this, several protocols have been described, but none of them has been shown to be more effective than other. Moreover, the total removal of cellular and nuclear components is crucial, as cellular or subcellular fragments left can trigger an immunological response. On the other hand, biological activity of the remaining ECM and preservation of the 3D architecture of the tissue, including a wide microvascular system, must be maintained ^(24–30)^.

Generally, two different approaches for cellular repopulation have been described: (i) infusion of cells via the conserved vascular network and (ii) direct injection of cells into the ECM ^(30–32)^. In the present study, we chose the second approach due to the significant reduction of the brain size after complete decellularization. Hence, we proposed a new method for decellularized brain scaffold production that could be used as a template for tissue bioengineering as the ECM structure is preserved.

## MATERIAL AND METHODS

### Experimental Model

Mice (*Mus musculus*, C57/Bl6 lineage) weighting 25-30 g, were obtained from the University’s Animal Facility (CEDEME/UNIFESP). All protocols were approved by the University’s Committee of Ethics in the Use of Animals (CEUA #2101180516). Every effort was made to minimize animal suffering and reduce the number of animals used. For the surgical proceedings, mice were anesthetized with Ketamine 75 mg/Kg and Xylazine 10 mg/Kg by intraperitoneal injection. Trichotomy and skin antisepsis were performed in abdominal site with chlorhexidine-alcohol and sterile cotton gauzes. Mice used to harvest the brain were placed in a surgical table at 37°C in dorsal decubitus.

### Decellularization process

Brains were obtained from adult C57BL/6 mice, placed inside a 2 mL tube and soaked in liquid nitrogen for 10 min. After this rapid freezing process, the brains were thawed at room temperature and washed with 10 mM PBS for 5 min. This freeze-thawing process was done 3 times. After this, each brain was soaked in a 10 mL solution of 1 % SDS, and agitated for 24 h. The decellularized brains were washed 3 times with 10 mM PBS for 5 min.

### Residual nucleic acid and protein quantitation

Nucleic acid quantitation was performed in samples of 100 mg of decellularized matrix or control fresh tissue, triturated in 1 mL Trizol (Life Technologies, USA) using a Polytron Pt 1200 Cl homogenizer (Kinematica AG, Switzerland). One mL of homogenate was transferred to propylene tubes and 250 μL of chloroform were added. After, tubes were sealed, homogenized and centrifuged 10000 × *g*; 15 min at 4°C. The supernatant was discarded and 300 μL of ethanol were added, and the mixture was gently homogenized in vortex and incubated at room temperature for 3 min. The solution was centrifuged 10000 × *g*; 5 min at room temperature. The DNA pellet was washed twice with 0.1 M citrate buffer and suspended in 1 mL of 75 % ethanol, incubated for 15 min with homogenization. The DNA solution was centrifuged at 10000 × *g*; 5 min at room temperature and the supernatant was gently removed, the pellet was dried for 5 min under vacuum and 100 μL of DNase free water were added.

DNA and RNA were isolated from native and decellularized tissue to perform the nucleic acid quantitation. The samples were triturated in 1 mL of Trizol using a Polytron Pt 1200 Cl homogenizer (Kinematica AG, Littau, Switzerland). One mL (1 mL) of homogenate was transferred to propylene tubes and 250 μL of chloroform were added. After, the tubes were sealed, homogenized and centrifuged 10,000xg, for 15 min at 4° C. The supernatant was discarded and 300 μL of ethanol were added, gently homogenized and incubated at room temperature for 2-3 min. Centrifuged at 10,000xg, for 5 min at room temperature. DNA pellet was washed twice with 0,1 M citrate buffer and suspended in 1 mL of 75 % ethanol, incubated for 15 min with periodic homogenization, centrifuged at 10,000 × *g*, for 5 min at room temperature and the supernatant was gently removed, dried for 5 min in vacuum and 100 μL of DNase free water were added. Nucleic acid content was quantified using NanoDrop ND-1000 (Thermo Fisher Scientific, USA). DNA and RNA contents were normalized to the initial dry weight of the samples. Base pair length of residual DNA was determined.

The total protein of the decellularized and non-decellularized brains was extracted using Mammalian Cell Extraction Kit (Biovision, USA) following the recommendations of the manufacturer, and after that the protein was quantified with Qubit (Thermo Fisher Scientific) fluorimeter.

### Characterization of decellularized tissues constituents

Decellularized and fresh tissues were fixed in 4 % paraformaldehyde and embedded in paraffin. Tissue sections (5 μm) were obtained and used for histological analysis. Haematoxylin and eosin (H&E) staining was performed according to standard protocol to evaluate the tissue morphology, specially the absence of residual cellular nuclei in the decellularized tissues, also confirmed by 4′,6-diamidino-2-phenylindole fluorescent dye (DAPI, 1:1000, Molecular Probes, USA) staining.

Basal membrane was evaluated by Periodic Acid-Schiff (PAS) staining, and Masson’s trichrome staining was used to stain collagen fibers. Alcian blue staining was used to evaluate the presence of sulfated glycosaminoglycans and picrosirius red was used to evaluate the presence of collagens type I, II, and III. All reagents used for the histological analysis were purchased from Sigma-Aldrich (USA), and the images were acquired with an Olympus BX61 microscope (USA).

### Culture of Neuro2a (mouse neuroblastoma)

Neuro2a cells were cultured according to the recommended protocol from ATCC (American Type Culture Collection, USA). Briefly, cells were cultured in a 75 cm^2^ culture bottle with 15 mL of culture medium composed of high glucose DMEM (Dulbecco’s Modified Eagle’s Medium, GibcoBRL, USA), 1 % penicillin / streptomycin (GibcoBRL), 1 % glutamine (Sigma-Aldrich) and 10 % fetal bovine serum (FBS; Cultilab, Brazil). Cells were kept in an incubator at 37° C and 5 % CO_2_. Upon reaching 60 % confluence cells were trypsinized and subcultured in the ratio of 1:5.

### Immunostaining

The control brains or decellularized brains were immunolabeled with anti-laminin antibody (Merck Millipore, USA), followed by incubation with fluorophore-conjugated secondary antibody (Alexa 488 or 594; Molecular Probes, USA) plus DAPI. Coverslips were assembled with Fluoromount G (Electron Microscopy Sciences) and then analyzed using an inverted confocal microscope (Leica Microsystems, Germany). Image overlays were generated using ImageJ software (NIH, USA).

Neuro2a cells were injected on decellularized brains and kept in an incubator at 37°C and 5 % CO_2_. After this period, cells were fixed with 4 % paraformaldehyde, permeabilized with 0.1 % Triton X-100 (Sigma-Aldrich), and immunollabeled with anti-beta III tubulin (Tubbeta3) (Merck Millipore) followed by incubation with fluorophore-conjugated secondary antibody (Alexa 488 or 594) plus DAPI. Coverslips were assembled with Fluoromount G (2: 1 in PBS) and then analyzed by an inverted confocal microscope. Image overlays were generated using ImageJ software.

### Repopulation of decellularized brain with Neuro2a cells

Repopulation of decellularized brain was performed after sterilization of the tissue with peracetic acid (PAA) 1:1000 in 20 % ethanol, placed on 24-well culture plate. After washing the samples 6 times with high glucose DMEM, 1 × 10^6^ Neuro2a cells were injected into the decellularized brain, and culture medium was added, until the incubation time periods were reached (24 and 72 h). After these periods of time, the repopulated decellularized brain was analyzed.

### Scanning electron microscopy

Samples of repopulated and non-repopulated decellularized brain fixed in 2.5 % glutaraldehyde solution buffered in 0.1 M sodium cacodylate solution, pH 7.2. After the fixation process, samples were washed with 0.1 M sodium cacodylate buffer, pH 7.2 and submitted to metallic impregnation. To make this, samples were incubated in 2 % osmium tetroxide in 0.1 M sodium cacodylate buffer, pH 7.2 for 2 h, washed with 0.1 M sodium cacodylate buffer pH 7.2 three times during 15 min, and incubated in 1 % tannic acid water solution for 45 min, followed by two washes in distilled water (10 min each). After metallic impregnation, samples were dehydrated gradually in 50-70-90 % ethanol (twice, 30 min each) and 100 % (three times, 30 min each) and then, samples were submitted to a drying process in a critical point chamber (Balzers CPD 030, Lichtenstein) using CO_2_. Samples were then coated with a thin layer of 20-30 nm thickness of gold (Sputtering, Leica Microsystems, Germany) and scanned on a FEI Quanta 250 FEG scanning electron microscope (ThermoFisher, USA).

### Statistical Analysis

The data were analyzed using the Statistical Package for Social Sciences (SPSS version 23.0, USA). The results were expressed by average ± SEM, and significance was tested by *t*-test or one-way ANOVA, as appropriate. The value of p < 0.05 was considered significant.

## RESULTS

### Murine brains were fully decellularized by SDS

After fast freezing in liquid nitrogen followed by 24 h in 1 % SDS, the mice brains were analyzed to investigate the remaining of cells, nucleic acids and proteins. Figure 1A presents a representative image showing that decellularized the brain became smaller, with less blood after the decellularization procedure. Further direct confocal microscopy analysis on a decellularized mouse brain section revealed the absence of nuclei stained with DAPI (Figure 1D), as compared to the clear presence of DAPI-stained nuclei in fresh non-decellularized mouse brain (Figure 1B). The absence of nuclei was confirmed by H&E staining (Figure 1C and E). Therefore, the fully decellularized murine brains were then named decellularized brain matrix (DBM).

**Figure 1:**
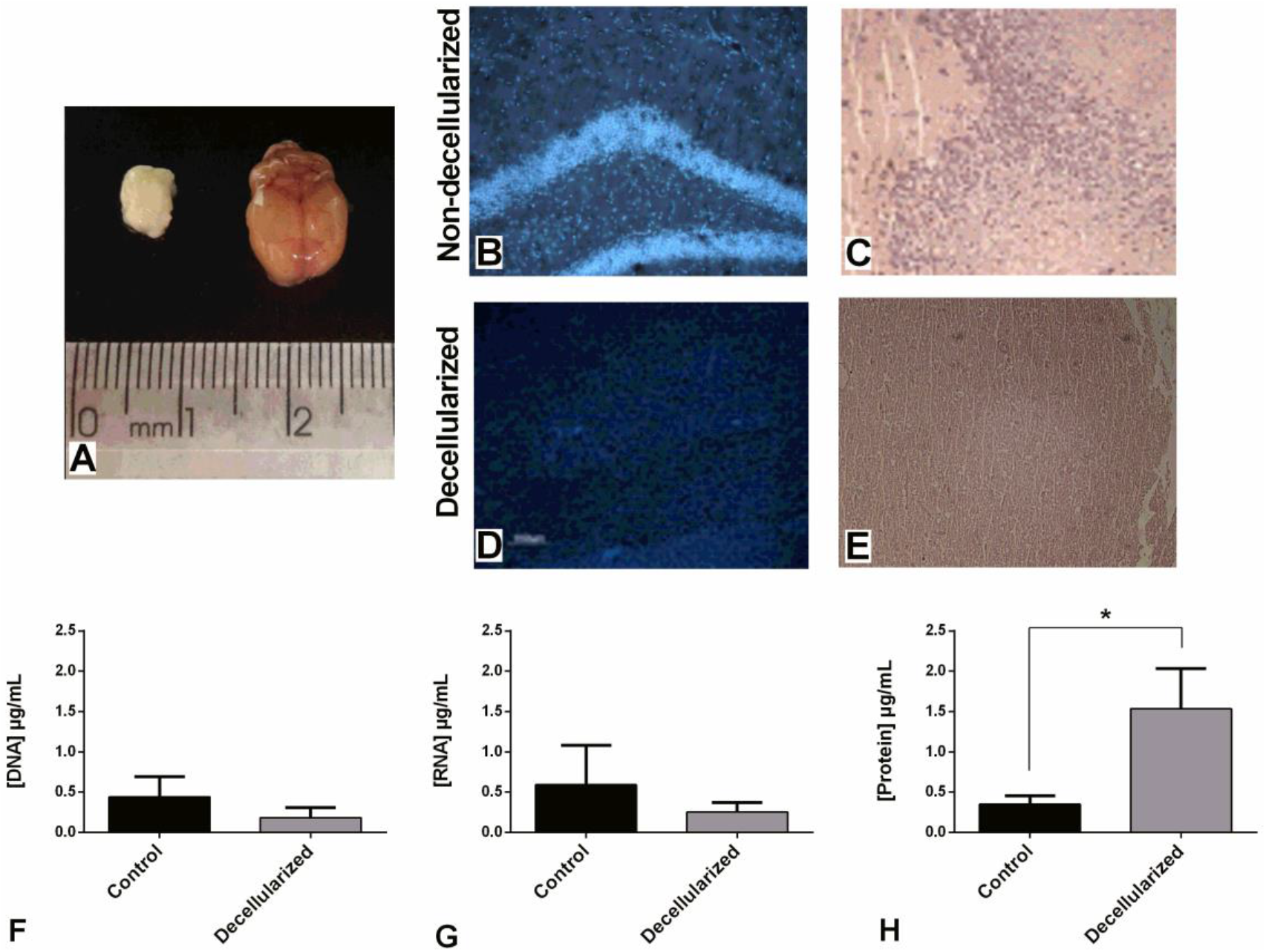
(A) Morphology change of the decellularized brain (left) and non-decellularized brain (right). Representative DAPI staining on 2 μm cryosections obtained from a non-decellularized (B) and a decellularized (D) mouse brain section. Representative haematoxylin-eosin (H&E) staining on 2 μm cryosections obtained from a non-decellularized (C) and a decellularized (E) mouse brain section. Scale = 200 μm. DNA (F) and RNA (G) quantification showed lower concentrations of nucleic acids in the decellularized scaffold compared with the non-decellularized tissue (Control), when both normalized with their dry weight. Protein quantification (H) showed higher concentrations of protein in the decellularized scaffold compared with the non-decellularized tissue (Control), when both normalized with their dry weight. T student test: *p < 0.05.

### Decellularization reduces the amount of nucleic acids leaving a higher protein concentration per weight of DBM

The quantification of DNA, RNA, and protein was performed in order to evaluate the effectiveness of decellularization. DNA, RNA, and protein concentrations were normalized by brain weight. The amount of DNA (Figure 1F) and RNA (Figure 1G) decreased after the decellularization process, whereas the the total protein concentrations was statistically higher after decelularization (Figure 1H, *p = 0.0155).

### DBM maintains essential ECM proteins after decellularization

Alcian Blue stains acidic polysaccharides such as glycosaminoglycans (Figures 2A and B). Masson’s Trichrom staining of collagen IV shows that its architecture was not affected by decellularization protocol (Figures 2C and D). Periodic Acid–Schiff (PAS) is a staining method used to detect polysaccharides such as glycogen, and glycoproteins, glycolipids and mucins in tissues. PAS staining showed that although the decellularization process was performed with a detergent (SDS), the basal lamina remained preserved (Figures 2E and F). Hematoxylin and eosin (H&E) staining is used to stain cell nuclei in tissues and to control reinforcing the effectiveness of decellularization process (Figures 2G and H).

**Figure 2:**
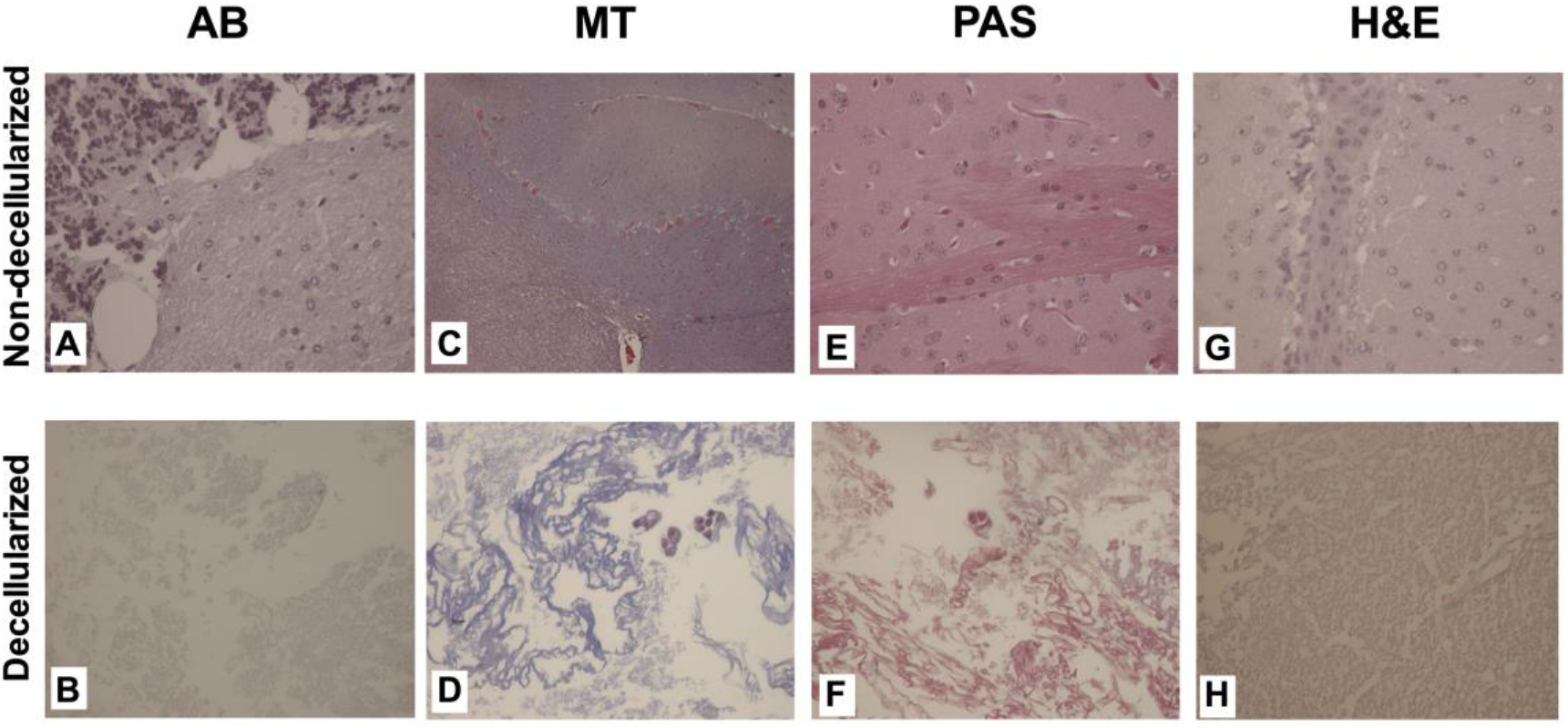
Representative Alcian Blue staining (A) and (B). Representative Masson’s Trichome (C) and (D). Representative PAS staining (E) and (F). Representative H&E staining (G) and (H). Cryosections obtained from a non-decellularized (Above) and a decellularized (Below) mouse brain section. Scale = 50 μm.

### DBM maintain collagen and laminin after decellularization

The immunofluorescence performed in non-decellularized brains and in DBM identified the presence of laminin, a protein crucial for the repopulation process. The results showed the presence of laminin (Figures 3A and 3B). Picrosirius red staining was performed to observe the presence of collagen type I, II and III (Figures 3C and 3D). Under polarized light, collagen bundles appear green, red, or yellow, and are easily differentiated from the black background ^33^. These two assays showed that laminin and collagen type I, II, and III were also preserved after the decellularization process.

**Figure 3:**
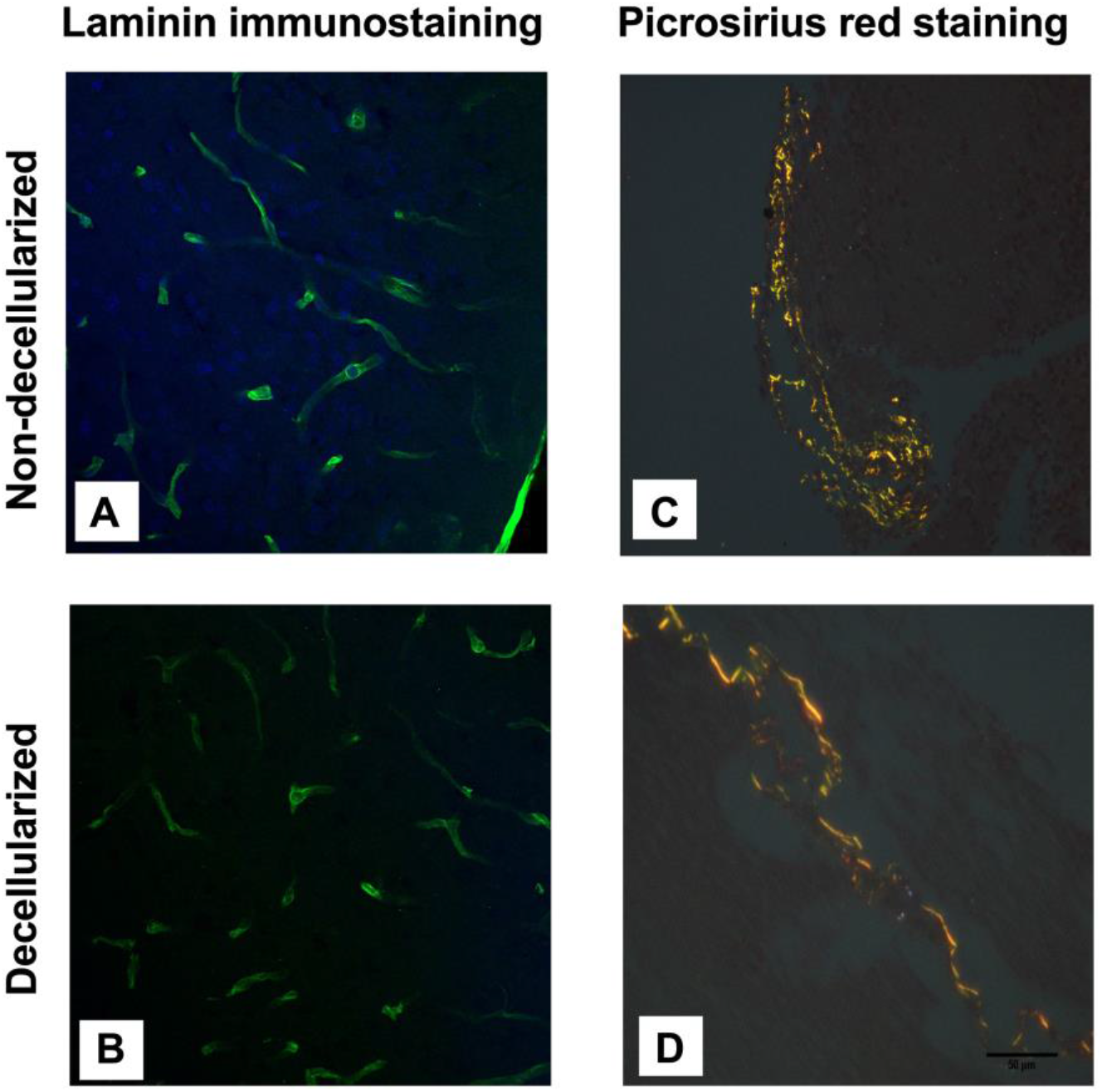
Representative Laminin immunostaining (green) with nuclear staining DAPI (blue) of non-decellularized (A) and decellularized (B) mouse brain section. Representative Picrosirius Red staining on 2 μm cryosections obtained from a non-decellularized (C) and a decellularized (D) mouse brain section. Scale = 50 μm.

### SEM microscopy shows the architecture of the DBM repopulated with Neuro2a cells

We also aimed to analyze if Neuro2a cells adhered when grown on DBM as they would do while cultured on plastic or glass. SEM analysis of slices of the internal region of DBM showed the presence of fibers of different lengths and thicknesses (Figure 4A) and holes that appear to be ripped cell nuclei (Figure 4B) from inner sections. In slices of the internal region of the DBM we could also observe the preservation of grooves (Figure 4E) while in the visualization of the external region of the DBM we could notice the presence of a vessel (Figure 4F). After recellularization for 24 h we observed that cell spreading and extended cellular protrusions were rare (Figure 4C and D), suggesting that there was no evidence of neuronal differentiation of Neuro2a cells, indicating that during this period of incubation the scaffold did not induce neuronal differentiation.

**Figure 4:**
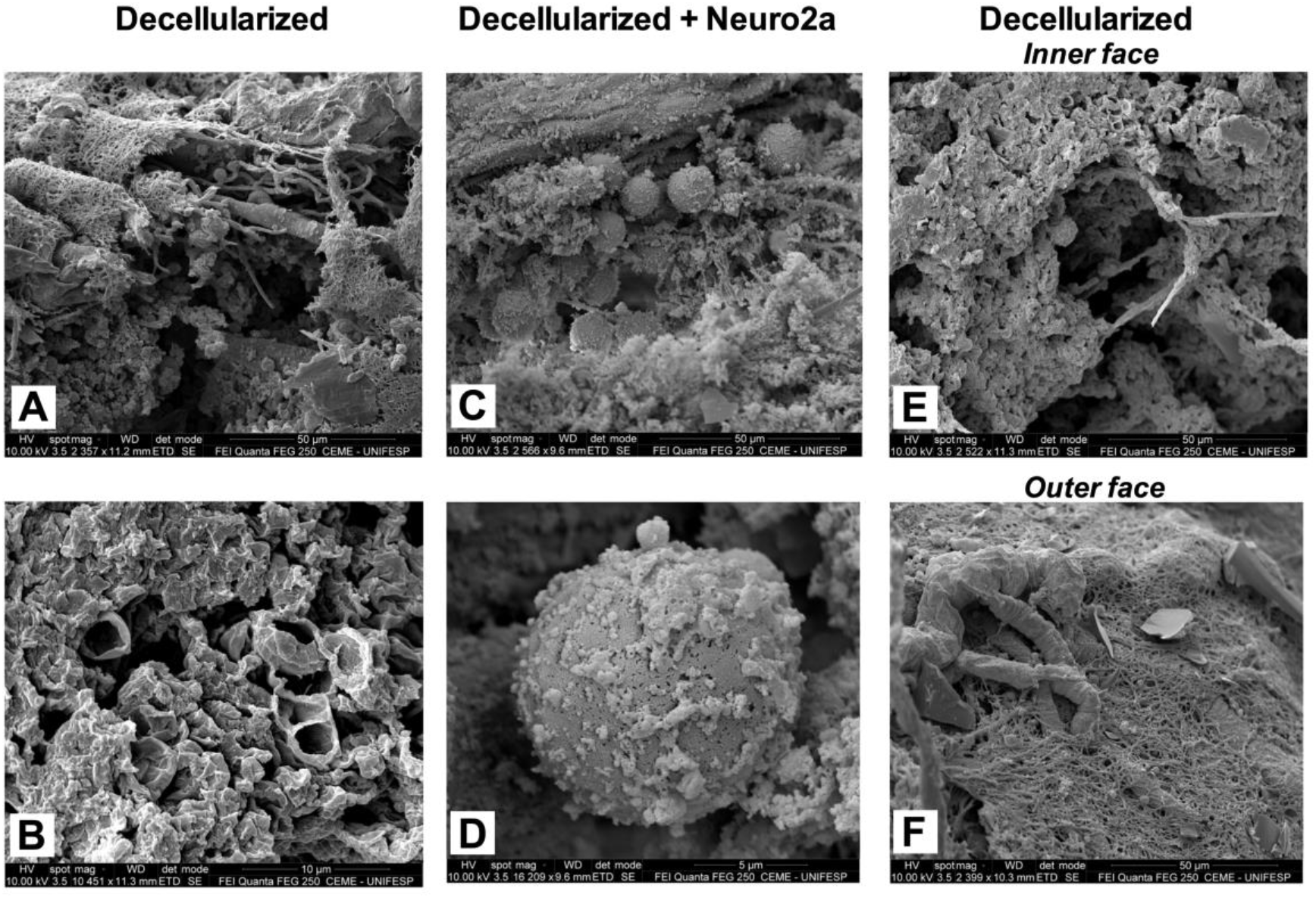
SEM images of DBM. (A) Internal region of DBM showing fibers of different lengths and thicknesses. (B) Internal region of DBM showing holes. Scale = 10 μm. Scale = 50 μm. (C) Internal region of DBM 24h after recellularization with Neuro2a cell. Scale = 50 μm. (D) Internal region of DBM 24h focusing on one cell. Scale = 10 μm. (E) Internal region of DBM showing the preservation of grooves. Scale = 50 μm. (F) External region of DBM showing the presence of a vessel. Scale = 50 μm.

### DBM maintained cells in an undifferentiated state

After injecting suspensions of Neuro2a cells into the DBM, they were incubated at 37°C and 5 % CO_2_ for 24 and 72 h. After fixing and performing the histological sections of the DBM, it was possible to find Neuro2a cells by immunostaining (Figure 5A and F), showing that the scaffold was able to maintain the cells alive and undifferentiated. There was no evidence of neuronal differentiation of Neuro2a cells (more evident TUBB3 marking and neurite extension), confirming that during these periods of incubations the scaffold did not induce neuronal differentiation, as observed by SEM (Figure 4).

**Figure 5:**
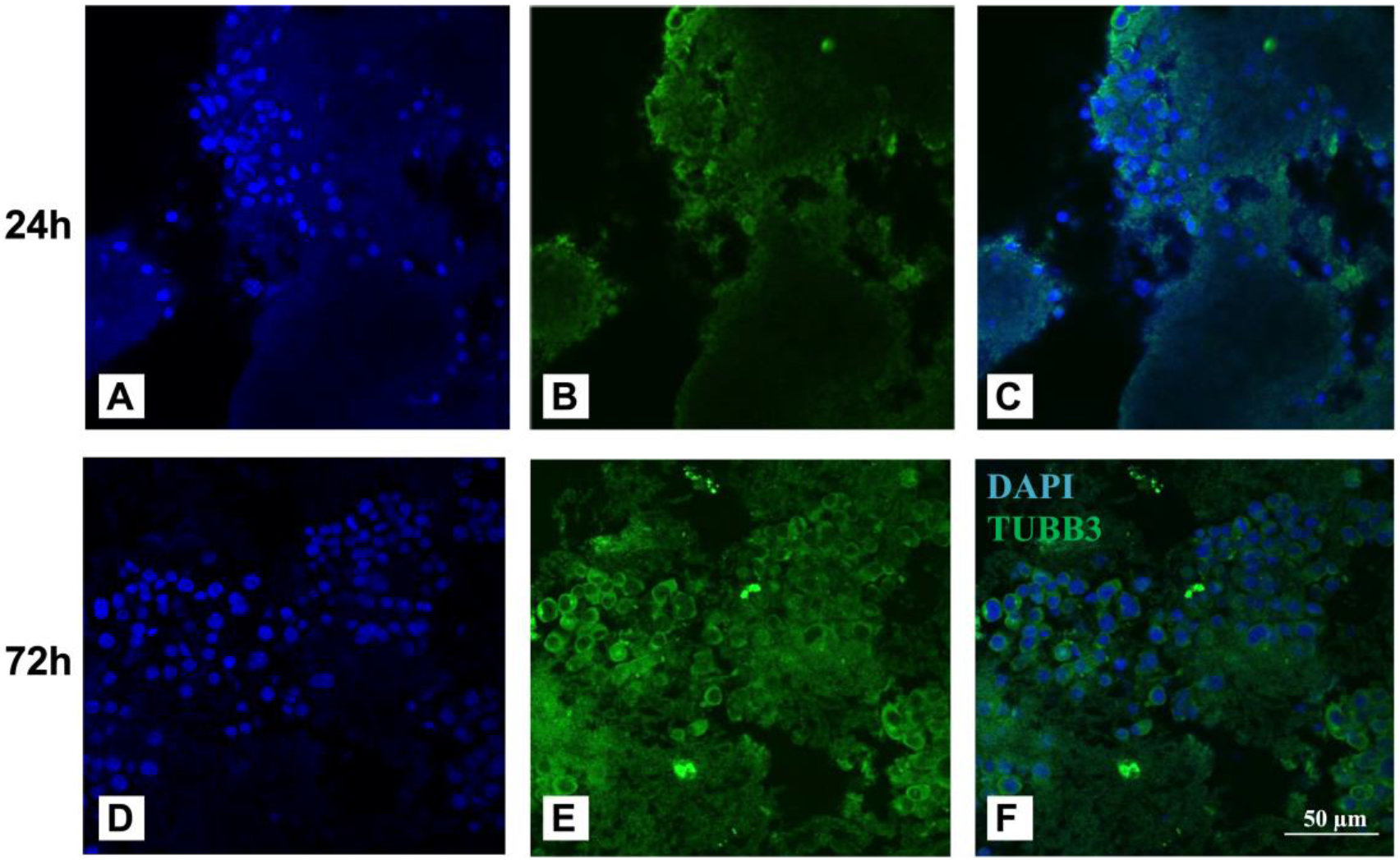
Neuro2a cells cultures inside DBM for 24h (A-C), 72h (D-F). Marker for mature neuron (Tubulin beta 3; TUBB3; green) (B,E), nuclear staining (DAPI; blue) (A,D) and merges of images (C,F). Scale = 50 μm.

## DISCUSSION

In the present work, we investigated the use of ECM scaffolds from murine brains (DBM) as a platform for repopulation of decellularized tissue. Our results showed a fast and straightforward method based on freezing and thawing murine brains followed by soaking them in a 1 % SDS solution for complete decellularization. This concentration was able to decellularize the brains with minimal residues of DNA reminiscent and preserving structures as the basement membrane, collagen IV, as well as adhesive proteins such as laminin. For the first time it was shown that a simple and rapid process involving just one chemical reagent could successfully decellularize murine brains.

The ECM has a crucial role in the development and repair of tissues. It offers support to cellular organization, and participates of cell growth and proliferation by the interaction with growth factors and cytokines ^(34–37)^. Therefore, the preservation of the ECM after the decellularization process showed here is very important to help cell reorganization after repopulation. In this work, after establishing a novel simple method for complete murine brain decellularization, and preservation of ECM structures, we also showed that the remaining scaffold (DBM) is biocompatible, being capable of maintaining Neuro2a cells alive for 72 h.

Few data regarding brain decellularization demonstrated complete decellularization with different methods ^38^. Decellularization regarding the regeneration of the CNS has been attempted by different techniques such as: detergent ^39^, detergent-enzimatic combination ^40^, enzymatic ^41^, and physical methods such as freezing and thawing and applying high hydrostatic pressure ^42^. Each of these protocols have their advantages and limitations. Notwithstanding, there are only a few studies concerning brain decellularization and its use for repopulation or as a scaffold.

Zhu et al. decellularized mice cerebellum and the resulting cerebellar scaffold retained neurosupportive proteins and growth factors and when tested with neural stem cells (NSCs) in vitro, was found to be cytocompatible ^43^. De Waele et al. also decellularized mice cerebellum and the survival and growth of seeded NSC cells was monitored for 1-7 weeks in culture, indicating the ability of the acellular brain sections to support sustained ex vivo growth of NSC retaining their NSC phenotype (undifferentiated) ^44^. These data corroborate with the results that we have described in this study, although we used a faster more straightfoward method for decellularization and fast-growing mouse neuroblastoma cell line (Neuro2a) with a shorter period of incubation (24 and 72 h) inside the decellularized brain scaffolds (DBM). It is worth mentioning that this study has some limitations and shall be continued analyzing the composition of the DBM in more depth.

## CONCLUSION

Decellularized scaffolds derived from an organ can be used as a platform to understand decellularization and cell repopulation. Although further studies are still required to validate the present data in translational models, our results are promising due to the fact that we could fully decellularize murine brains with a fast (24 h for complete decellularization) and simple (only one detergent and 3 freezing and thawing processes) method. The decellularization process also retained essential ECM proteins that are crucial for cell adhesion, migration, proliferation and consequently repopulation of the tissue. Lastly, the decellularized brain scaffold showed to be biocompatible, as neural cells were injected inside their structure and could be identified by immunohistochemistry.

## AKNOWLEDGEMENTS

This work was supported by Fundação de Amparo à Pesquisa do Estado de São Paulo – FAPESP (2012/00652-5), Conselho Nacional de Desenvolvimento Científico e Tecnológico – CNPq (402319/2013-3; 465656/2014-5), and Coordenação de Aperfeiçoamento de Pessoal de Nível Superior – CAPES (Finance Code 001).

