## Supplemental text for "A novel decellularization method to produce brain scaffolds"

### Supplementary Material

*Electrophysiology*

Cells were cultured for 72h with complete medium on culture dishes (Indifferentiated cells) and on DBM (Indifferentiated cells on DBM). Cells were also cultured with neuronal differention medium (The same complete medium with decreased FBS concentration from 10% to 0.5% and with the addition of 10 mM retinoic acid). After 72h the cells were trypsinized using trypsin (Cultilab) in 10 mM PBS. After this cells where plated on culture dishes and left to adhere for 24h before being used for electrophysiology. Recording glass pipets were pulled from borosilicate glass (World Precision Instruments Inc., Berlin, Germany), using a two-stage puller (Sutter P-0; Sutter Instruments, Novato, CA). Pipet tips were fire-polished using a flame polisher (MF-83; Narishige, Tokyo, Japan). Whole-cell voltage-clamp recordings were carried out at room temperature. The compositions of extracellular and intracellular buffers were: extracellular buffer -125 mM NaCl, 25 mM $NaHCO_3$, 1.25 mM, $NaH_2PO_4$, 3 mM KCl, 2 mM $CaCl_2$, 1 mM $MgCl_2$, 25 mM glucose and 3 mM pyruvic acid; intracellular buffer - 135 mM potassium gluconate, 7 mM NaCl, 10 mM HEPES, 2 mM $Na_2ATP$, 0.3 mM $Na_2GTP$ and 2 mM $MgCl_2$. The cells were patched with borosilicate glass electrodes (resistance 6 - 10 MΩ) and the measurements performed using an Axopatch Amplifier 200A (Molecular Devices Corp., Union City, CA). Current responses were elicited by step depolarizations from a holding potential of -70 mV to 60 mV, for 50 ms each pulse, in intervals of 10 mV. Data were analyzed using Clampfit (Molecular Devices) and GraphPad Prism version 6.0.

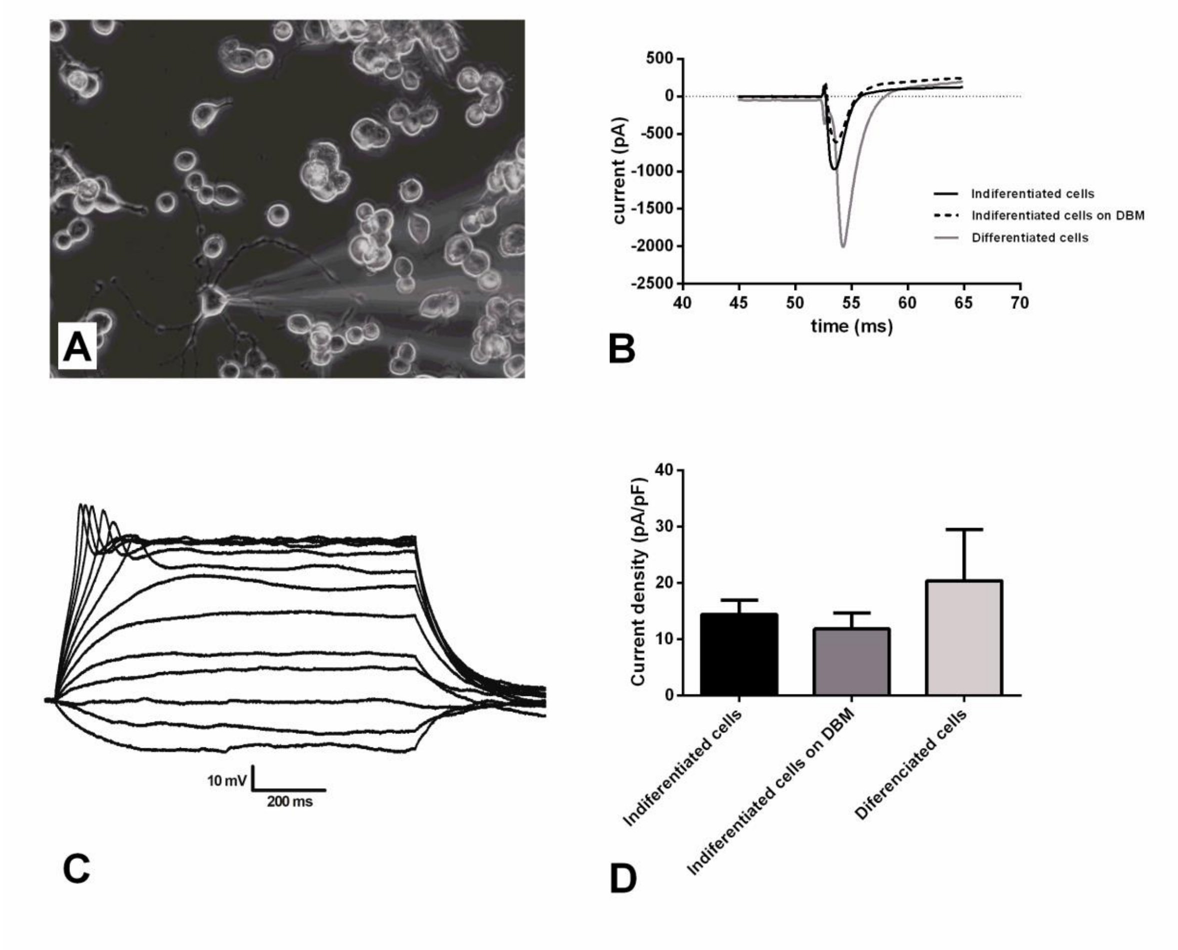


**Supplementary Figure 1:** (A) Illustrative image of a 72h differentiated Neuro2a cell being clamped with the recording glass pippete. Scale = 30 µm. (B) Representative action potentials of a differentiated cell evoked by current application. (C) Representative current traces obtained by voltage clamp of Neuro2a cells. (D) Analysis of the current density (pA/pF) of Neuro2a cells.
